# The study of lumbar ligamentum flavum hypertrophy induced in bipedal mice

**DOI:** 10.1101/723239

**Authors:** Zhenyu Zheng, Lei Qian, Xiang Ao, Peng Li, Yongxing Peng, Jun Chu, Tao Jiang, Zhongmin Zhang, Liang Wang

## Abstract

Lumbar spinal stenosis (LSS) is a common degenerative disease among the elderly. The role that mechanical stress-induced hypertrophic ligamentum flavum (HLF) plays in patients with LSS remains unclear. Here, we used a finite element analysis to investigate the stress characteristics on the ligamentum flavum (LF) and evaluate the feasibility of a mouse model of HLF. First, we induced a bipedal posture in mice by taking advantage of their hydrophobia. A micro-CT scan was performed to examine their spinal change during bipedal posture. A finite element analysis showed that the stress and strain on the upright posture were significantly increased compared with those on the sprawling posture. Tissue staining showed that the degeneration degree of the LF in bipedal standing group gradually increased over the modeling period. The amount of elastic fibers decreased under HLF, whereas the amount of collagen fibers, the number of the LF cells, and the expression of fibrosis-related factors increased. Compared with aged group, LF degeneration was more severe in the bipedal standing group. Our findings demonstrate that the increased stress caused by a posture change causes HLF and that a bipedal mouse model can be used to study HLF in vivo.

## Introduction

Lumbar spinal stenosis is one of the most common spinal degenerative diseases, and it has a high incidence among elderly individuals. LSS is usually caused by HLF (Costandi et al., 2015; Jirathanathornnukul et al., 2016; Sakai et al., 2017; Szpalski and Gunzburg, 2003). A degenerative and thickening of LF can cause spinal stenosis and compress the nerve roots or cauda equina, thereby leading to back pain and intermittent claudication (Park et al., 2001). A variety of factors including age, activity level, genetic composition, and mechanical stress accelerate the development of HLF (Fukuyama et al., 1995; Sairyo et al., 2005). To date, most researchers agree that an abnormal stress level can accelerate the degradation and hypertrophy of LF (Kong et al., 2009; Moon et al., 2012; Shafaq et al., 2012). Mechanical stress causes microinjuries to the LF tissue. Repeated microinjuries lead to chronic inflammation and subsequent tissue scarring, which eventually cause HLF (Kosaka et al., 2007; Sairyo et al., 2007; Schrader et al., 1999; Yoshida et al., 1992). In vitro studies have demonstrated that mechanical tension promotes collagen synthesis via the TGF-β1 pathway (Nakatani et al., 2002). However, no in vivo studies have directly demonstrated that mechanical stress induces LF hypertrophy.

In addition, because it is not necessary to remove the facet joints of patients with LSS during surgical decompression, it is difficult to observe the pathological relationship between HLF and facet joint degeneration. Although many studies have revealed the underlying molecular mechanisms in cells, the effects of abnormal stress and fibrosis-related molecules on HLF cannot be studied in vivo. Mice are a common and cost-efficient animal model for medical research. However, only one complicated stretch device has been used to model HLF in these animals (Saito et al., 2017), and its utility is limited by its complicated operation, and requirement for repeated anesthesia. Therefore, the effectiveness of this model has been questioned.

With the increased application of finite element analysis in medicine, soft tissue finite element research has gradually become available. The use of this method can help obtain the mechanical parameters that are difficult to capture using the traditional biomechanical method. Micro-CT combined with finite element research can help to intuitively analyze the mechanical problems at the microscopic level and enable the biomechanical analysis of the mouse LF. Based on a previous study of spinal degeneration (Ao et al., 2019), we used bipedal standing mice to make HLF. Finite element analysis was used to verify the hypothesis that the mouse LF is hypertrophied under abnormal tension. Furthermore, a histopathological analysis was employed to verify the validity of the model and thereby validate this model as a new mouse model for LF degeneration.

## Results

### Finite element analysis

In the sprawling and bipedal positions, the stress and strain on the LF are primarily concentrated at the junction between the LF and the facet joint. The von Mises stress was 1.79×10^−5^ MPa, and the maximum principal strain was 8.62×10^−5^ (Fig. 1A, C). The anteriorly tilting of the L5 in an upright standing posture can result in a pulling-force on the junction between the lower articular process of the L5 and the LF. When the LF is stretched, the stress and strain on the LF at the junction with the facet joint is remarkably increased, the von Mises stress value is 8.85×10^−2^ MPa, and the maximum principal strain is 6.64×10^−1^ (Fig. 1B, D).

**Fig. 1.**
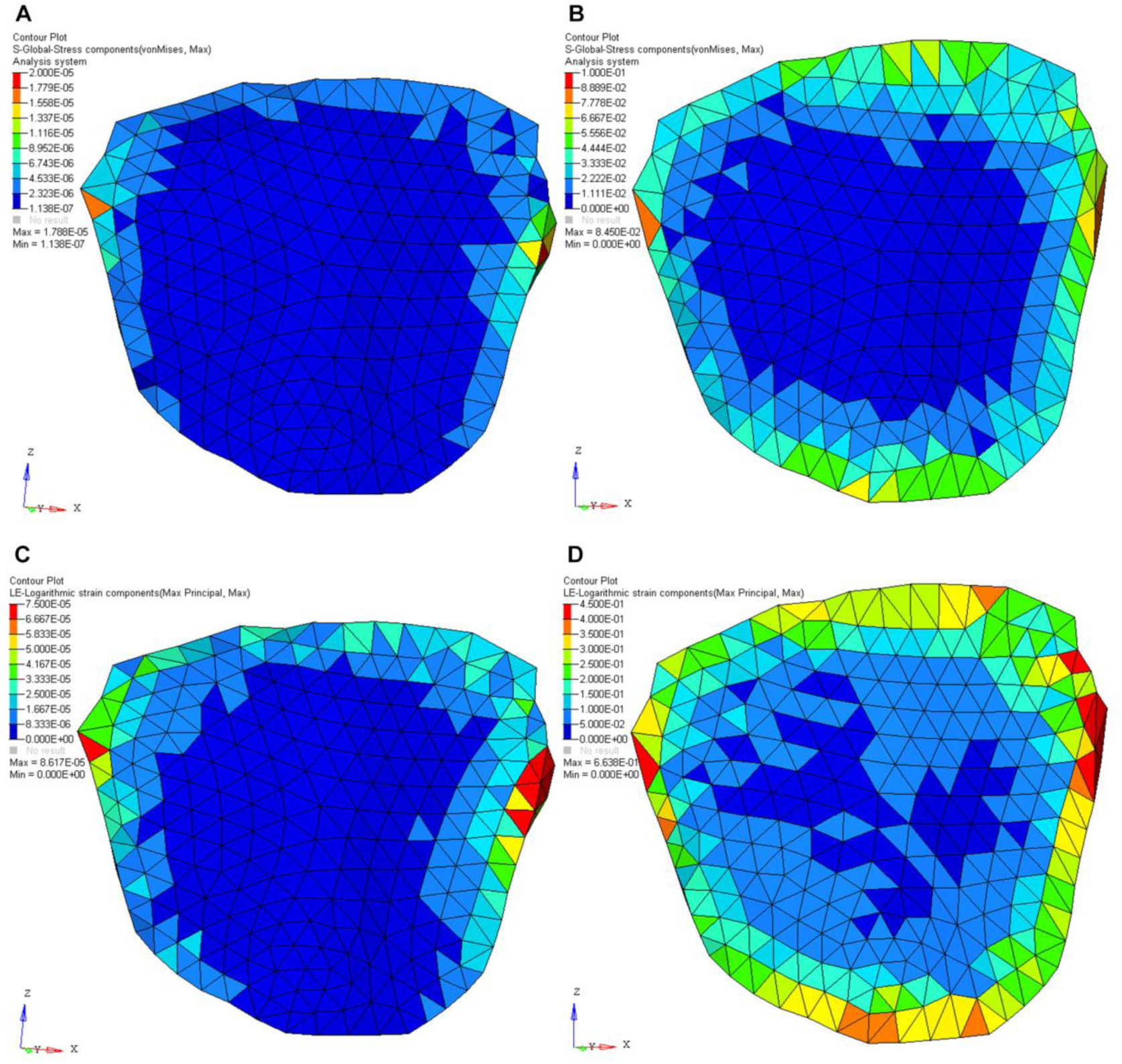
Changes in posture alter the stress on the mouse LF. The distribution and von Mises stress values of the LF in the sprawling (A) and upright standing positions (B); The distribution and values of the maximum principal strain of the LF in the sprawling (C) and the upright standing positions (D).

### Abnormal stress causes LF hypertrophy

H&E staining showed that the area of the LF in the bipedal standing group was larger than that in the control group (Fig. 3A, B). Furthermore, we observed that the area of the LF was larger in the 10-week bipedal standing group than that in the control and 18-month-old groups (Fig. 3C, D). Although ageing can also cause HLF, No significant difference was observed between the 18-month-old group and the control group (p = 0.06; Fig. 3D).

**Fig. 2.**
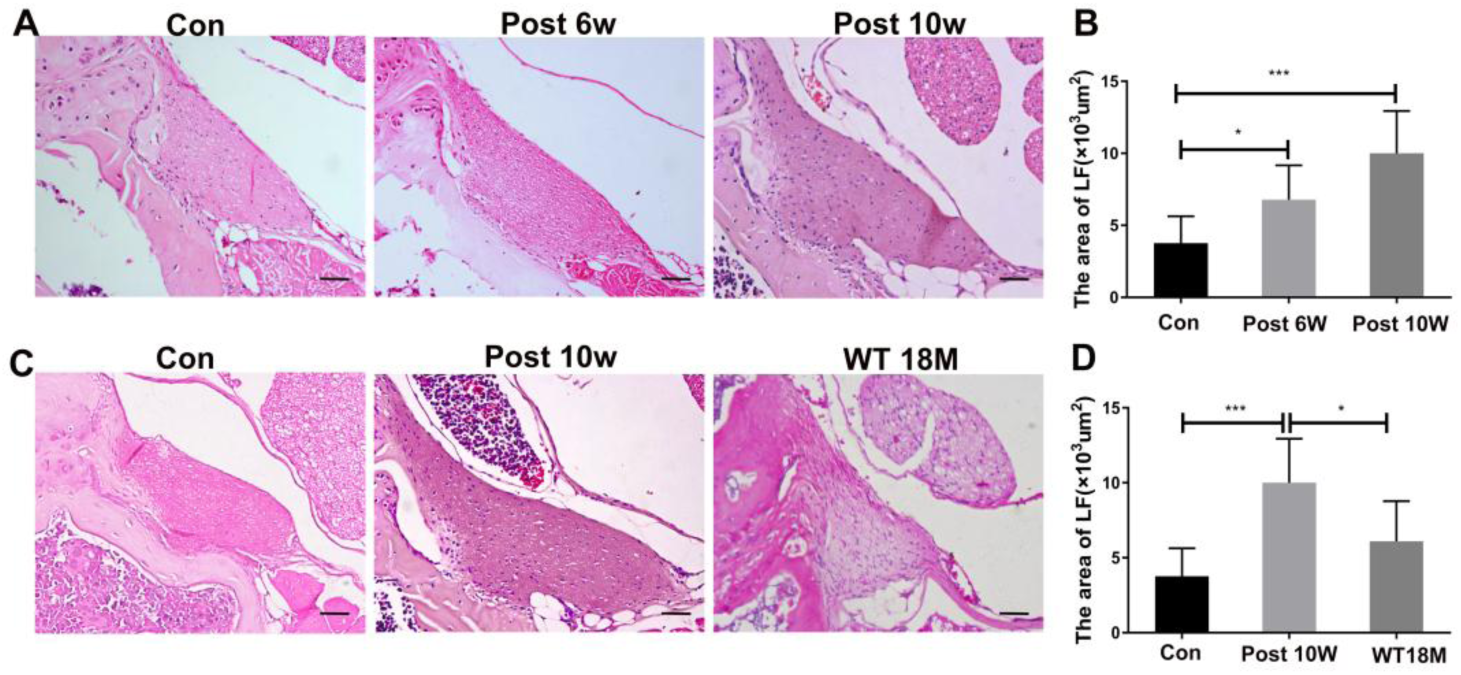
Mechanical stress is the main causes of HLF. (A) H&E staining of LF from different groups. (B) The statistical results of mice LF area in different groups, (n = 8). (C) H&E staining of LF specimens from different groups. (D) Statistical results of the mice LF area in different groups, (n = 8). ANOVA was used. P-value cut-off of 0.05 was used, *p < 0.05, ***p < 0.001. Error bars show means ± SD, scale bars = 20 µm; Con = control group; Post 6W = post 6-week bipedal standing group; Post 10W = post 10-week bipedal standing group; WT 18M = wild-type aged mice at 18 months.

**Fig. 3.**
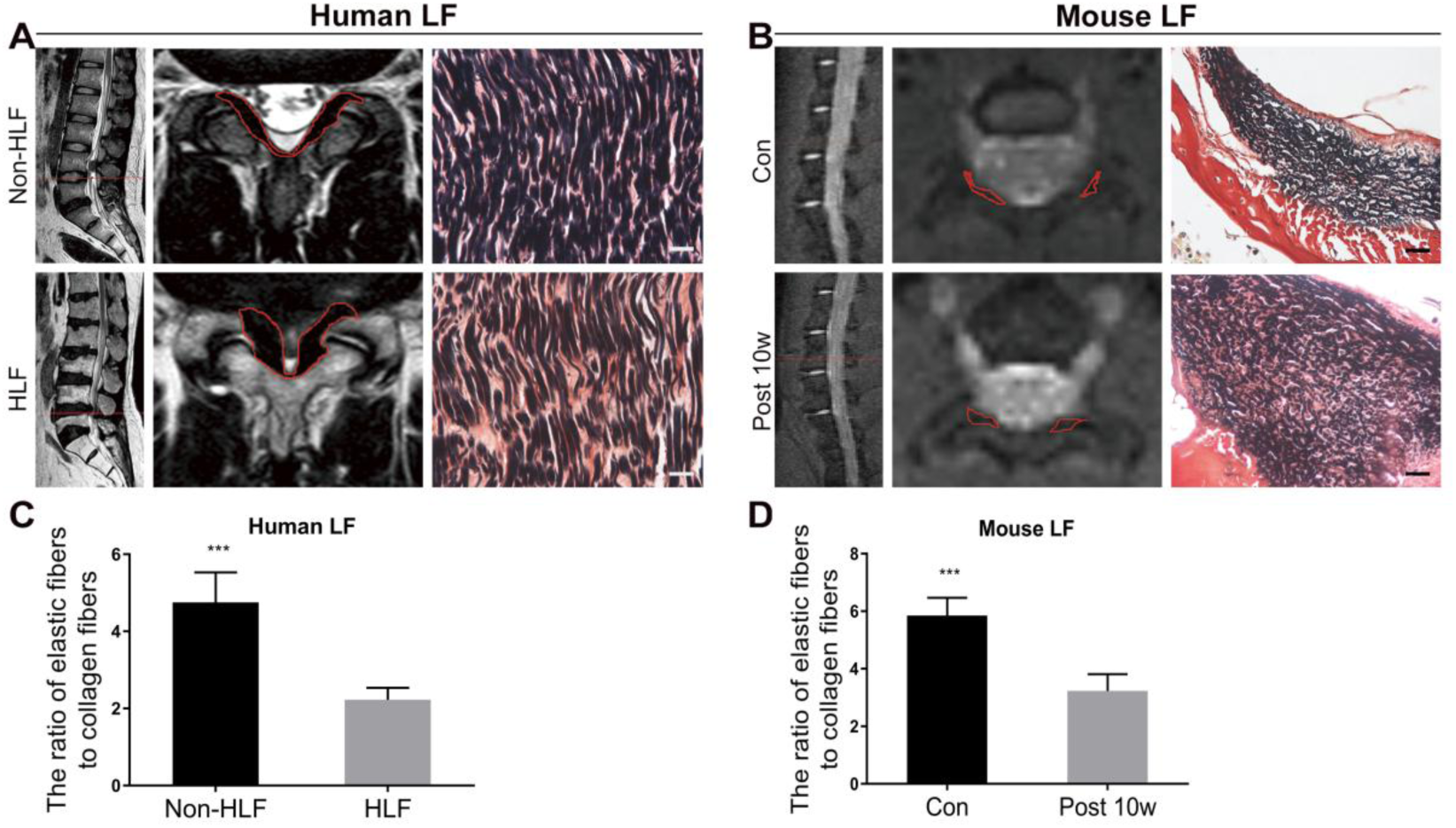
Pathological changes of the LF in a bipedal mice model simulate HLF in humans. (A) VVG staining of LF specimens and lumbar magnetic resonance imaging (MRI) of patients with spinal canal stenosis and nonspinal stenosis; the red line indicates the contour of the LF. In the HLF group, VVG staining shows that the purple-black elastic fibers are significantly reduced, but the red collagen fibers are significantly increased; (B) VVG staining of LF specimens and lumbar MRI of mouse models and control groups; the red line indicates the contour of the LF; VVG staining of the LF specimens; (C) The ratio of elastic fiber to collagen fiber in the two groups of human LF, (n=8); (D) The ratio of elastic fiber to collagen fiber in the two groups of mouse LF, (n=8). ***p<0.001 (scale bars=10 µm; HLF=hypertrophied ligamentum flavum; Con=control group; Post 10w=10-week modeling group).

### Histological comparison between human and mouse LF degeneration

We compared the pathological changes to the LF in the mouse model with those of human LF specimen. In the Verhoeff-Van Gieson (VVG) staining of the human LF, the non-HLF was dominated by dense and neatly arranged elastic fiber bundles (black strip fibers) with a small amount of collagen fibers (red); The HLF had a decreased content of elastic fiber that was thinned, broken, and irregularly arranged. The proliferation and accumulation of collagen fibers was increased (Fig. 3A). The elastic-fiber-to-collagen-fiber-area ratio in the HLF was reduced relative to that in the non-HLF (Fig. 3C). Previous results are consistent with our findings (Kosaka et al., 2007). The pathological changes of LF in our mouse model were similar to those in the human samples. More elastic fibers with a small amount of collagen fibers were present in the LF in the control group. In the 10-week modelling group, the area of collagen fibers increased significantly (Fig. 3B). Although the area of the elastic fibers was also increased, the ratio of elastic fibers to collagen fibers decreased (Fig. 3D). These results indicate that the HLF pathology in the bipedal standing model is histologically identical to that in humans.

### Increased number of LF cells and the upregulation of fibrosis biomarker in the LF

During the process of HLF, in addition to changes in the extracellular matrix (ECM), the number of cells and cell types also change. An important feature of fibrosis is the conversion of fibroblasts to myofibroblasts that express alpha-smooth muscle actin (a-SMA) and secretory ECM (Wynn, 2008). We examined the changes in cell density and the number of a-SMA positive cells in the human and mouse LF samples respectively. In the human LF specimens, the density of the LF cells in the HLF cells was higher than that in the non-HLF cells (Fig. 4A and Fig. 4B). The number of a-SMA positive cells in the HLF cells was significantly increased compared with that in the non-HLF cells (Fig. 4E and Fig. 4F). Similarly, in the mouse model, the cell density in the LF of the bipedal standing 10-week group was significantly higher than that in the control group (Fig. 4C and 4D). Furthermore, the number of a-SMA positive cells in the LF of the bipedal standing 10-week group was higher than that in the control group (Fig. 4G and Fig. 4H).

**Fig. 4.**
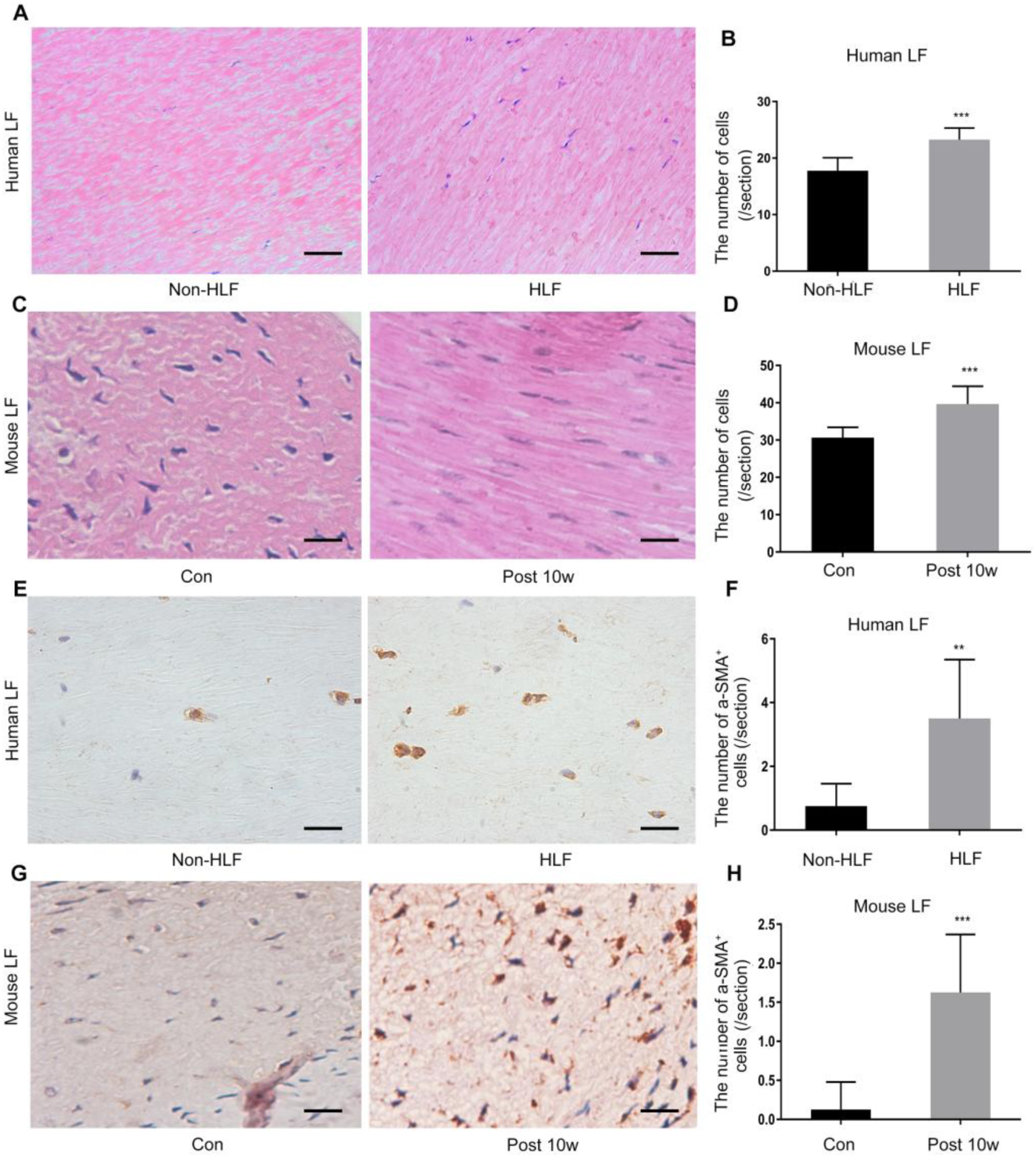
Changes in the cell density and number of a-SMA positive cells in the HLF and non-HLF cells in mice and humans. (A, B) The cell density in the human HLF was significantly increased; (n=8); (C, D) the cell density in the mouse LF was significantly higher in the bipedal standing 10-week group than in the control group; (n=8); (E, F) the number of a-SMA positive cells in the HLF of humans was significantly increased; (n=8); (G, H) The number of a-SMA positive cells in the LF of the bipedal standing group was higher than that in the control group; **p<0.01, ***p<0.001; scale bars (A): 50 μm; (E): 20 μm; (C, G): 10 μm.

### Increased expression of fibrosis-related factors in the bipedal HLF mouse model

To study the changes in the fibrosis-related factors in the LF of the bipedal mouse group, we harvested the LF tissue and evaluated the gene expression levels of collagen I, collagen III, a-SMA, MMP2, interleukin-1β and COX2 in the LF of the 10-week model and control groups using RT-qPCR. The results showed that all of these indicators were significantly higher in the LF in the 10-week group than that in the control group (Fig. 5 A-F).

**Fig. 5.**
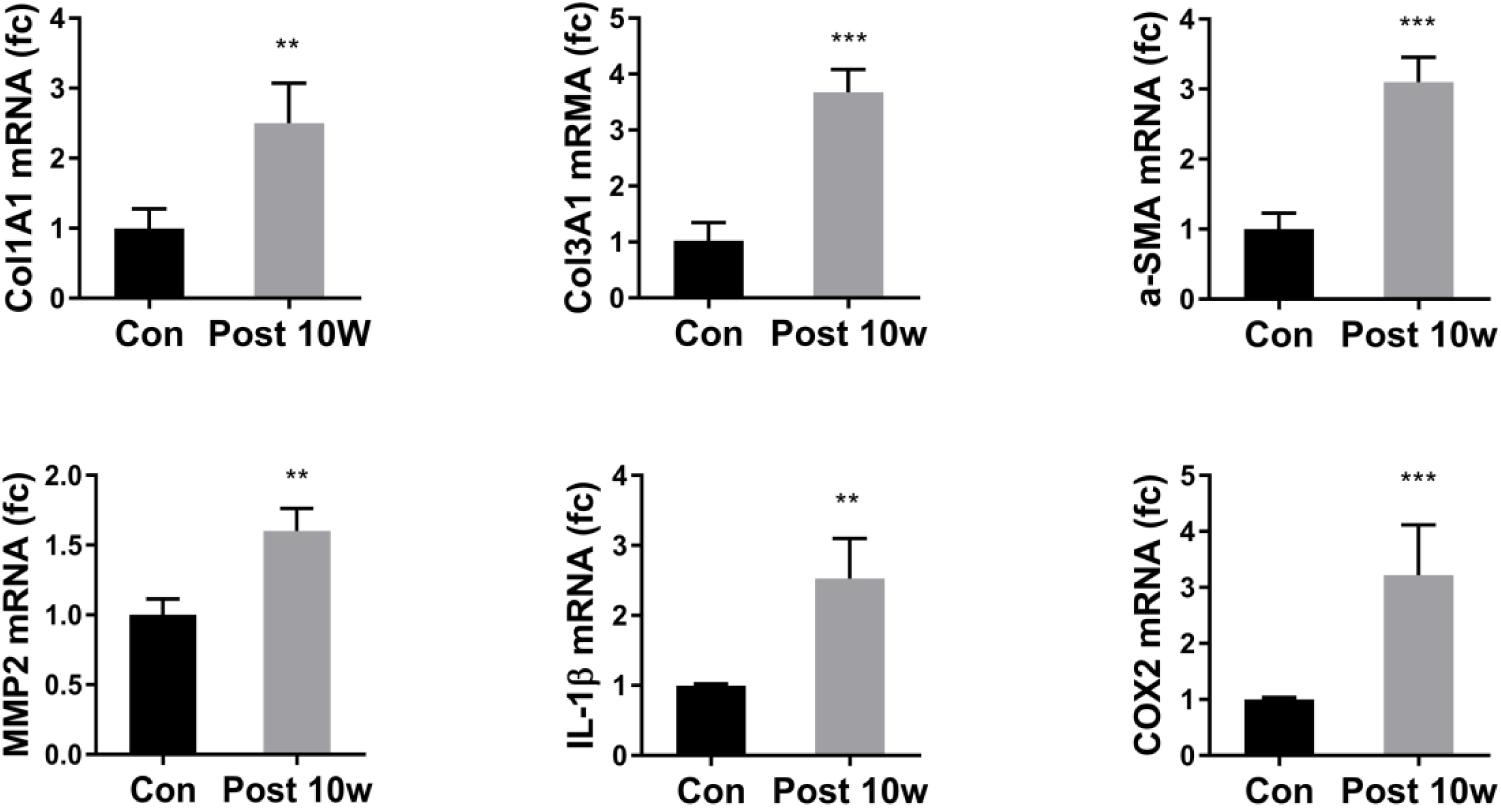
Increased mRNA expression of fibrosis-related factors in the bipedal mouse model. RT-qPCR was used to detect the expressions of COL1A1, COL3A1, a-SMA, MMP2, IL-1β and COX2 in the LF of the control and 10-week modeling groups (n=3); **p < 0.01, ***p <0.001.

## Discussion

Mechanical stress might control the metabolism of the LF matrix and the expression of various fibrosis-associated factors and angiogenic factors by affecting the biological behavior of LF cells (Hur et al., 2015; Nakatani et al., 2002). However, because of the lack of effective upright standing vertebrate models, most current experiments have focused on the study of cells, and in vivo research has rarely been reported. In a previous study, we found that bipedal standing can cause mouse spinal degeneration. Because the LF is an important structure in the posterior column, it is necessary to use this model to further study the pathological changes of the LF to provide an animal model for in vivo studies of HLF. The finite element analysis confirmed that the stress distribution within the mouse spine changed when the mouse was upright. This change might lead to the HLF. During the early stages of the model, the degenerative changes were first caused by the load-sharing effect of the facet joints. The mouse LF originates from the medial edge of the facet joint and is primarily located in this region. The axial sections showed that the LF appears as a triangular-like shape here. However, the LF is thinner at other locations and appears stringy. Therefore, abnormal stress to the facet joints might cause similarly abnormal stress on the LF. The prolonged modeling time is related to the increased degree of HLF. The results of the finite element analysis support this hypothesis and show that the L5 vertebral body flexed anteriorly in reference to the L6 when the mice were in bipedal standing positions. The stress and strain were primarily concentrated in the LF near the facet joint, which is at the main anatomical location of the LF. The finite element analysis confirmed that the relative position of the L5/L6 changes in the bipedal mouse model caused the lower L5 articular process to pull the LF, increasing the stress and strain on the LF and, ultimately, HLF.

This mouse model effectively simulates the pathological changes of human HLF. The normal LF in humans is an elastic structure composed of 80% elastic fibers and 20% collagen fibers (Viejo-Fuertes et al., 1998). Studies have reported that increased collagen fibers, broken elastic fibers, and increased cell numbers (Hayashi et al., 2017; Sairyo et al., 2005; Shafaq et al., 2012) in human HLF are associated with the increased expression of fibrosis-related factors (Lakemeier et al., 2013). Our bipedal mouse model confirmed the presence of increased collagen fibers, cell proliferation, and the expression of fibrosis-related factors in the LF. This finding is consistent with human LF pathology. In addition, the immunohistochemistry results showed that the number of a-SMA positive cells in human HLF was higher than that in the normal LF. Similarly, the number of a-SMA positive cells in the bipedal mouse model of HLF was also increased.

Facet joint degeneration is an important indicator of spinal degeneration, and it is associated with age and LF degeneration (Yoshiiwa et al., 2016). Our study shows that the area of the LF was significantly greater in the bipedal modeling group than in the aged group. The current model is reliable and simple. The results of the finite element analysis show that the stress on the LF is consistent with the histological changes. Compared with other types of models, the current model is easy to generate and exerts minimal trauma on the mouse. Compared with Takeyuki Saito et al. who examined the effect of loaded stress on LF biodynamics using an in vitro tension device loading model, repeated anesthesia is not necessary for the bipedal standing model. Furthermore, the modeling device is simple. The simple operation of the model effectively changes the sprawling posture into an upright posture among vertebrate animals, which changes the way that the spine is stressed. Based on this model, the effects of various cytokine levels on HLF and its mechanisms can be studied in vivo.

This study has some limitations. A long-term study is needed to understand the mechanisms that underlie HLF. In summary, we demonstrated for the first time that abnormal stress directly causes LF hypertrophy in a mouse model. The bipedal mouse model can be used to simulate human HLF pathogenesis, and this work provides a reference for in vivo studies of HLF.

## Materials and methods

### Animal experiments

A total of 32 eight-week-old C57BL/6 male mice (Guangdong Medical Laboratory Animal Center) were randomly assigned to control and experimental groups. The mice in the experimental group were placed in a cylindrical molding device (15 cm in diameter and 20 cm in height) that contained 5-mm-deep water at the bottom. The mice were maintained in a bipedal standing posture for 6 hours a day with a 2-hour break in the middle. The control mice were also placed in the modeling device without water at the bottom. After 6 and 10 weeks of modeling period, 8 mice were randomly chosen for euthanasia. Histological staining was performed to assess the degree of LF degeneration in eight mice. For quantitative reverse-transcription polymerase chain reaction (qRT-PCR) assay, another 32 eight-week-old C57BL/6 male mice were used. The Animal Experimental Ethics Committee of the Southern Medical University approved the animal experimental protocol based on the Animal Experiment Guide. A total of 16 male C57BL/6 mice were randomly sacrificed at the ages of 12 or 18 months to serve as an aged mouse model of LF degeneration. The degeneration degrees of the LF were evaluated.

### Finite element analysis

The mice were anesthetized and subjected to a micro-CT scan (LaTheta LCT-100S; Aloka, Tokyo, Japan). The mice were secured in the scanning capsule in the sprawling and bipedal standing postures. The scanning parameters were as follows: 55 kV, 109 μA, slice thickness 96 μm, exposure time 200 ms, and pixels 512×512. A total of 650 and 606 images of the L5-L6 segment were obtained in the DICOM data format with the mice in the sprawling and bipedal standing positions, respectively. The image data were imported to Mimics 14.0 (Materialise Corp., Leuven, Belgium) for 3D reconstruction (Fig. 6), and the reconstructed .stl files were imported into Hypermesh 13.0 (Altair, Troy, MI, USA) for meshing. Then, the data were introduced to Abaqus 6.14 (Dassault Systemes Simulia Corp., Providence, RI, USA) for a material property assignment. Both the sprawling and bipedal standing posture models included the vertebral cortical bone, the vertebral cancellous bone, the endplate, the annulus fibrosus, the nucleus pulposus, the ligament, and the joint capsule. The endplate is a shell unit, and the ligament (except the LF) and joint capsule are spring units. The LF is a truss unit, and the other components are tetrahedral units. The friction coefficient of the joint was 0.1 (Polikeit et al., 2003), and the material properties of each component are shown in Table 1.

**Table 1.** Composition and material properties of each component.

| Components | Material and property | Reference |
| --- | --- | --- |
| Annulus ground substance | Hyperelastic Mooney-Rivlin<br>$C_{01}=C_{10}=0.1$; $C_{02}=C_{11}=C_{20}=0.01$;<br>$D_1=D_2=35$ | (Lotz, 2005) |
| Nucleus pulposus | Hyperelastic Mooney-Rivlin<br>$C_{01}=0.01$; $C_{10}=0$; $D_1=100$ | (Lotz, 2005) |
| Endplate | Elastic, $E=100$, $\nu=0.2$ | (Shirazi-Adl, 1996) |
| Cancellous bone | Elastic, $E=50$, $\nu=0.2$ | (Hayes, 1997) |
| Cortical bone | Elastic, $E=148000$, $\nu=0.3$ | (Floor M. Lambers, 2011) |
| LF | Hyperelastic Mooney-Rivlin<br>E=15 (<6.2%), 19.5 (>6.2%) 0.3 | (Chen-Sheng<br>Chen,<br>2001) |

**Fig. 6.**
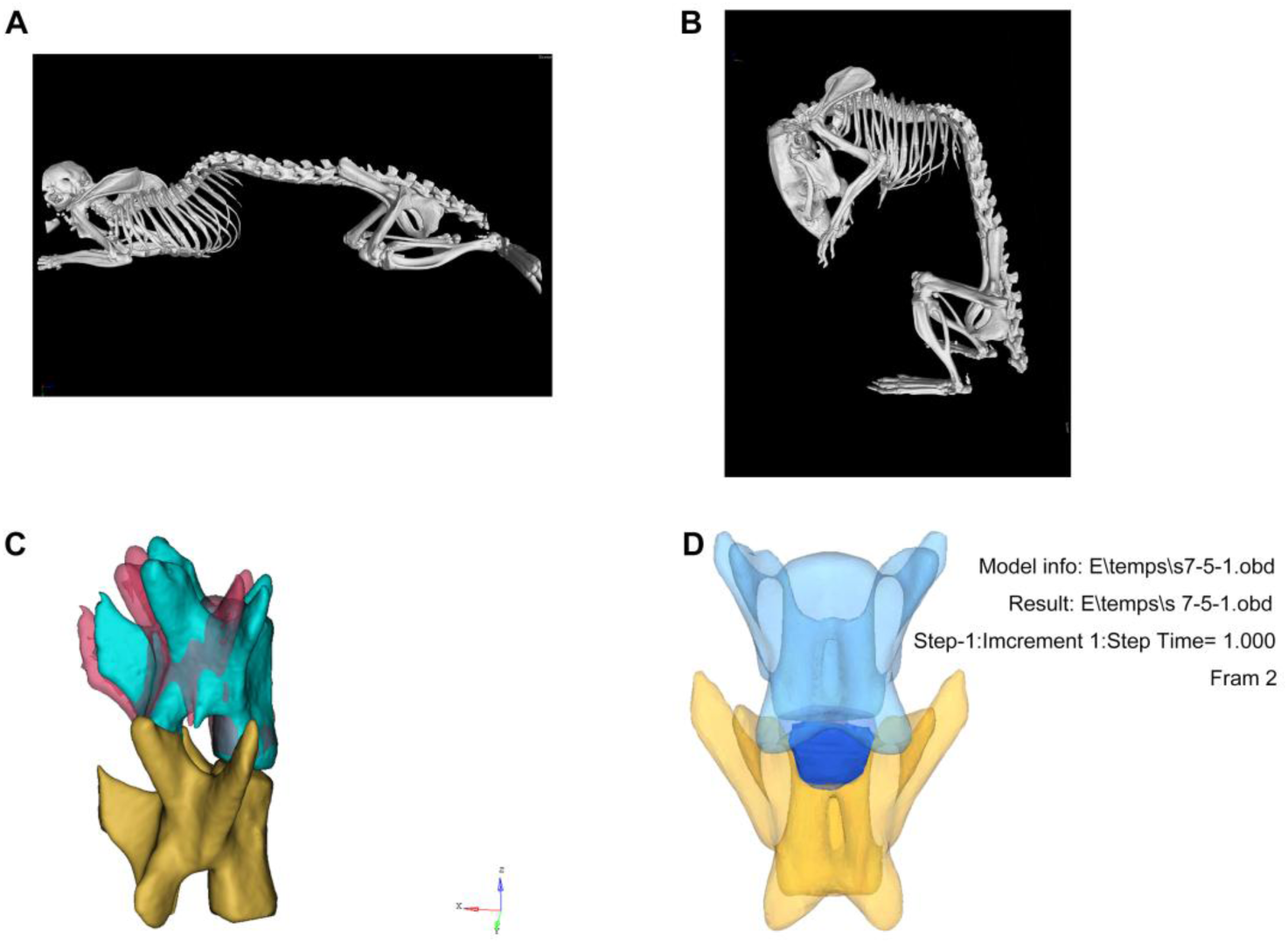
3D reconstruction of the mouse lumbar vertebrae. A 3D reconstruction model with using DICOM data for a mouse in the sprawling posture (A) and bipedal standing position (B); the relative position of the L5 after normalization in the sprawling and bipedal standing positions; the red area represents the L5 position in the sprawling posture, and the green area represents the L5 position in the upright standing position (C); the blue area represents the location of the LF (D).

To analyze the stress on the LF in mice in the sprawling posture, the vertebral body was secured at the ventral side, and a single force of 0.2 N (half the body weight of the mouse) was applied to the dorsal side of the vertebral body to calculate the degree and distribution of the stress and strain on the LF. For bipedal standing mice, the bottom of the L6 was secured, and a single force of 0.2 N was applied to the top of the L5 to calculate the degree and distribution of the stress and strain on the LF.

### Histological analysis

After the animals were euthanized, the intact L5, and L6 vertebrae were harvested and fixed in neutral formaldehyde. The specimens were embedded in paraffin after decalcification in an ethylenediaminetetraacetic acid solution and dehydration; Then, they were sectioned to a thickness of 4 µm. H&E staining was performed to observe the extent of the LF degeneration. Both human and mouse LF tissues were stained with VVG staining to assess the extent of LF fibrosis, and the staining procedure was performed according to the manufacturer’s instructions.

### Immunohistochemistry

The sections were deparaffinized and incubated in a citrate buffer and heated for 16 hours at 60°C. The samples were incubated with 3% H_2_O_2_ for 10 minutes to quench the endogenous peroxidase activity. The sections were washed three times for 5 minutes in phosphate-buffered saline (PBS) and blocked in 5% goat serum at room temperature for 1 hour. The sections were stained with primary antibodies anti-aSMA (A11111, ABclonal, Wuhan, China). After incubation at 4°C overnight, the sections were washed three times for 5 minutes in PBS. Then, the sections were incubated with a species-matched HRP‐labelled secondary antibody (1:500) at 37°C for 1 hour. DAB was used as the chromogen, and hematoxylin was used as a counterstain. After mounting, the corresponding protein expression was observed under a microscope.

### Human LF samples

The Ethics Committee of the Third Affiliated Hospital of Southern Medical University approved the experiment, and each patient provided informed consent before surgery. During surgery, specimens were obtained from eight patients diagnosed with LSS and HLF with a mean age of 63.4 ± 4.0 years and eight patients with lumbar disc herniations but without HLF with a mean age of 58.7 ± 3.5 years. The dorsal layer of LF tissues were taken from the same anatomical region (L4/5). The differences in the age and gender distributions of the two groups were not significant.

### Image quantification

The thickness of the human LF was measured at the facet joint of the L4/5 level with axial T1-weighed MRI (Philips, Amsterdam, Netherlands) as previously described (Yu Zhang, 2010). The mouse LF was observed at the facet joint of the L5/6 level on the axial T1-weighed MRI (PharmaScan, Bruker, Germany). The images for the histological and immunohistochemical analyses were obtained using a digital light microscope (Axio Scope.A1, Zeiss, Jena, Germany). ImageJ (National Institutes of Health) was used to calculate the area of interest. The area was calculated three times, and the average value was taken. The investigators (X.A., Y.P., and P.L.) were blind to the group assignment.

### qRT-PCR analysis

RNeasy Micro Kit (74004, Qiagen, Germany) was used to extract total RNA from LF cells. The purity and concentration of the total RNA was then measured using a NanoVue Plus spectrophotometer. HiScript II Q SuperMix was used for the reverse transcription reaction. The qPCR reaction was performed with ChamQ SYBR qPCR Master Mix. The primers that were used in the experiment were synthesized by Sangon Biotech (Shanghai, China). β-actin was used as the internal reference. The data were analyzed using the 2^-ΔΔCt^ method (Livak and Schmittgen, 2001). The primers used for the real-time reactions were as follows: (1) COL1A1 forward primer (5’-GCTCCTCTTAGGGGCCACT-3’) and reverse primer (5’-CCACGTCTCACCATTGGGG-3’); (2) COL3A1 forward primer (5’-TAAAGAAGTCTCTGAAGCTGATGG-3’) and reverse primer (5’-ATCTATGATGGGTAGTCTCATTGC-3’); (3) MMP2 forward primer (5’-CCCTCAAGAAGATGCAGAAGTTC-3’) and reverse primer (5’-TCTTGGCTTCCGCATGGT-3’); (4) COX2 forward primer (5’- AACCGAGTCGTTCTGCCAAT-3’) and reverse primer (5’-CTAGGGAGGGGACTGCTCAT-3’); (5) IL-1β forward primer (5’-GGG CTGGACTGTTTCTAATGCCTT-3’) and reverse primer (5’-CCATCAGAGGC AAGGAGGAAAACA-3’); (6) a-SMA forward primer: (5’-CCACCATCTGCC TGAAATCC-3’) and reverse primer: (5’-GCTTCTTGTCCAGCCTCCTC-3’); and (7) β-actin forward primer (5’-CTTAGTTGCGTTACACCCTTTCTTG-3’) and reverse primer (5’-CTGCTGTCACCTTCACCGTTCC-3’).

### Statistical analyses

All statistical analyses were performed using SPSS 22.0 (SPSS Inc, Chicago, IL, USA). The data were expressed as means±SD. A t-test was used to analyze data with a normal distribution and homogeneity of variance. Welch’s t-test was used to analyze data that were not normally distributed and showed heterogeneity of variance. One-way analysis of variance (ANOVA) was used for comparisons among multiple groups. P-values<0.05 indicated a significant difference.

